# Super-enhancer profiling reveals ThPOK/ZBTB7B, a CD4^+^ cell lineage commitment factor, as a master regulator that restricts breast cancer cells to a luminal non-migratory phenotype

**DOI:** 10.1101/2024.09.21.614267

**Authors:** Camila D Arcuschin, Kamin Kahrizi, Rosalyn W Sayaman, Carolina DiBenedetto, Yizhuo Shen, Pedro J Salaberry, Ons Zakroui, Cecilia Schwarzer, Alessandro Scapozza, Paola Betancur, Julie D Saba, Jean-Philippe Coppé, Mary-Helen Barcellos-Hoff, Dietmar Kappes, Laura van ‘t Veer, Ignacio E Schor, Denise P Muñoz

## Abstract

Despite efforts to understand breast cancer biology, metastatic disease remains a clinical challenge. Identifying suppressors of breast cancer progression and mechanisms of transition to more invasive phenotypes could provide game changing therapeutic opportunities. Transcriptional deregulation is central to all malignancies, highlighted by the extensive reprogramming of regulatory elements that underlie oncogenic programs. Among these, super-enhancers (SEs) stand out due to their enrichment in genes controlling cancer hallmarks. To reveal novel breast cancer dependencies, we integrated the analysis of the SE landscape with master regulator activity inference for a series of breast cancer cell lines. As a result, we identified **T**-**h**elper-inducing Poxviruses and Zinc-finger (**PO**Z)/**K**rüppel-like factor (ThPOK, *ZBTB7B*), a CD4^+^ cell lineage commitment factor, as a breast cancer master regulator that is recurrently associated with a SE. ThPOK expression is highest in luminal breast cancer but is significantly reduced in the basal subtype. Manipulation of ThPOK levels in cell lines shows that its repressive function restricts breast cancer cells to an epithelial phenotype by suppressing the expression of genes involved in the epithelial-mesenchymal transition (EMT), WNT/β-catenin target genes, and the pro-metastatic TGFβ pathway. Our study reveals ThPOK as a master transcription factor that restricts the acquisition of metastatic features in breast cancer cells.

## Introduction

In recent years, breast cancer mortality has declined due to broader screening, early detection, better classification, and the discovery of new targeted therapies. However, metastatic breast cancer remains incurable, and many patients lack effective treatments (1, 2).

Breast cancer is classified based on the expression of different receptors: estrogen and/or progesterone receptors (ER/PR+), amplification or overexpression of epidermal growth factor 2 (HER2+), or the absence of these three markers in the case of triple negative breast cancer (TNBC). However, this classification is complemented by the molecular intrinsic subtyping based on transcriptional profiles that form the PAM50 signature and divides them in luminal A (Lum A), luminal B (Lum B), Her2-enriched, basal-like and normal-like (3, 4).

Genetic and epigenetic factors contribute to the heterogeneity of breast cancer, with aberrant epigenetic changes driving cancer progression. Super-enhancers (SEs) are emerging as key regulatory regions in tumorigenesis, aiding in diagnosis and classification, understanding subclonal evolution, and predicting drug responses (5, 6). They are characterized by unusually high signals of histone modifications such as H3K27ac and H3K4me1, and chromatin regulatory proteins such as BRD4, the mediator complex, and cell-type specific transcription factors (TF) (7, 8). SEs control the expression of genes essential for normal cells’ identity, and in cancer cells regulate genes involved in survival, oncogenesis and drug resistance (7-12). This suggests that identifying SEs and their associated genes can reveal previously unappreciated cell dependencies and vulnerabilities, as well as inform about cancer biology, clinical diagnosis and therapeutic guidance (13). In breast cancer cells (BCCs), SEs regulate genes involved in immune evasion (14) and therapeutic resistance (15, 16), and their perturbation by bromodomain and extraterminal (BET) protein inhibition restrains proliferation and promotes apoptosis (12, 17).

ThPOK (**T**-**h**elper-inducing Poxviruses and Zinc-finger (**PO**Z)/**K**rüppel-like factor), encoded by the *ZBTB7B* gene, is a transcription factor that belongs to the POK family of transcriptional repressors containing BTB/POZ domains. It was originally known for regulating extracellular matrix (ECM) genes in fibroblasts (18, 19) and CD4 T lymphocyte lineage commitment (20), but its role in other contexts is less known. While other members of the POK family have been implicated in carcinogenesis (21), there is no such oncogenic function described for ThPOK under non-manipulated circumstances. However, transgenic mice constitutively expressing ThPOK in T cells (T-cell specific *ThPOK* transgene, *ThPOK*^const^ mice) develop thymic lymphomas resembling human T-cell acute lymphoblastic leukemia (T-ALL) (22). Only recently, ThPOK has been implicated in regulating lipid synthesis and lipid droplet secretion by the lactating murine mammary gland (23).

Here, we reveal a novel role for ThPOK as a master regulator in BCCs. The ThPOK/*ZBTB7B* gene is associated with a SE in cells derived from different breast tumor subtypes, a feature that distinguishes key transcriptional regulators in cancer. ThPOK expression levels are higher in breast tumors compared to normal adjacent tissues, with the highest levels observed in cells/tumors of the luminal subtype. Conversely, a major repressed status is observed in TNBC cell lines or tumors, likely driven by specific CpG methylated sites. Manipulation of ThPOK levels revealed a role for ThPOK in limiting BCCs plasticity to an epithelial non-migratory phenotype by restricting the expression of ECM genes, epithelial-mesenchymal transition (EMT) and stemness factors, and WNT and TGFβ pathways.

## RESULTS

### Ubiquitous super-enhancers associate with key regulators of gene expression in breast cancer cell lines

Since SEs control oncogenes and regulators of cellular identity (5, 7, 24), we reasoned that characterizing SE profiles in both normal and cancer cells can reveal cancer-specific dependencies. To map the regulatory landscape of breast cancer, we used chromatin immunoprecipitation followed by sequencing (ChIP-seq) to determine the genome-wide distribution of H3K27ac in BCC lines derived from primary tumors. We combined our data with available H3K27ac ChIP-seq datasets from different BCC lines. After quality control of 80 input/H3K27ac ChIP-seq datasets, we analyzed 37 samples from 17 breast cancer cell lines (**Table S1**), identifying H3K27ac peaks and SEs using MACS and ROSE2 algorithms (11, 25). The consolidated set includes 5917 SEs (**Fig. 1a**) proximal (+/-10Kbp) to 9911 genes. Our analysis confirmed that the 200 most variable SE regions (**Table S2)** cluster the cell lines according to their subtypes with few exceptions (**Figs. S1a-b**). Functional analysis of the complete SE set using the Genomic Regions Enrichment of Annotations Tool (GREAT) (26) showed SE-associated genes enriched in terms related to cell differentiation, mRNA metabolism, cell signaling and cell architecture (**Fig. S1c** and **Table S3**). In summary, using this strategy, we obtained a consolidated SE set that reflects known grouping of the cell lines and at the same time is associated with genes with functions relevant to carcinogenesis.

**Figure 1.**
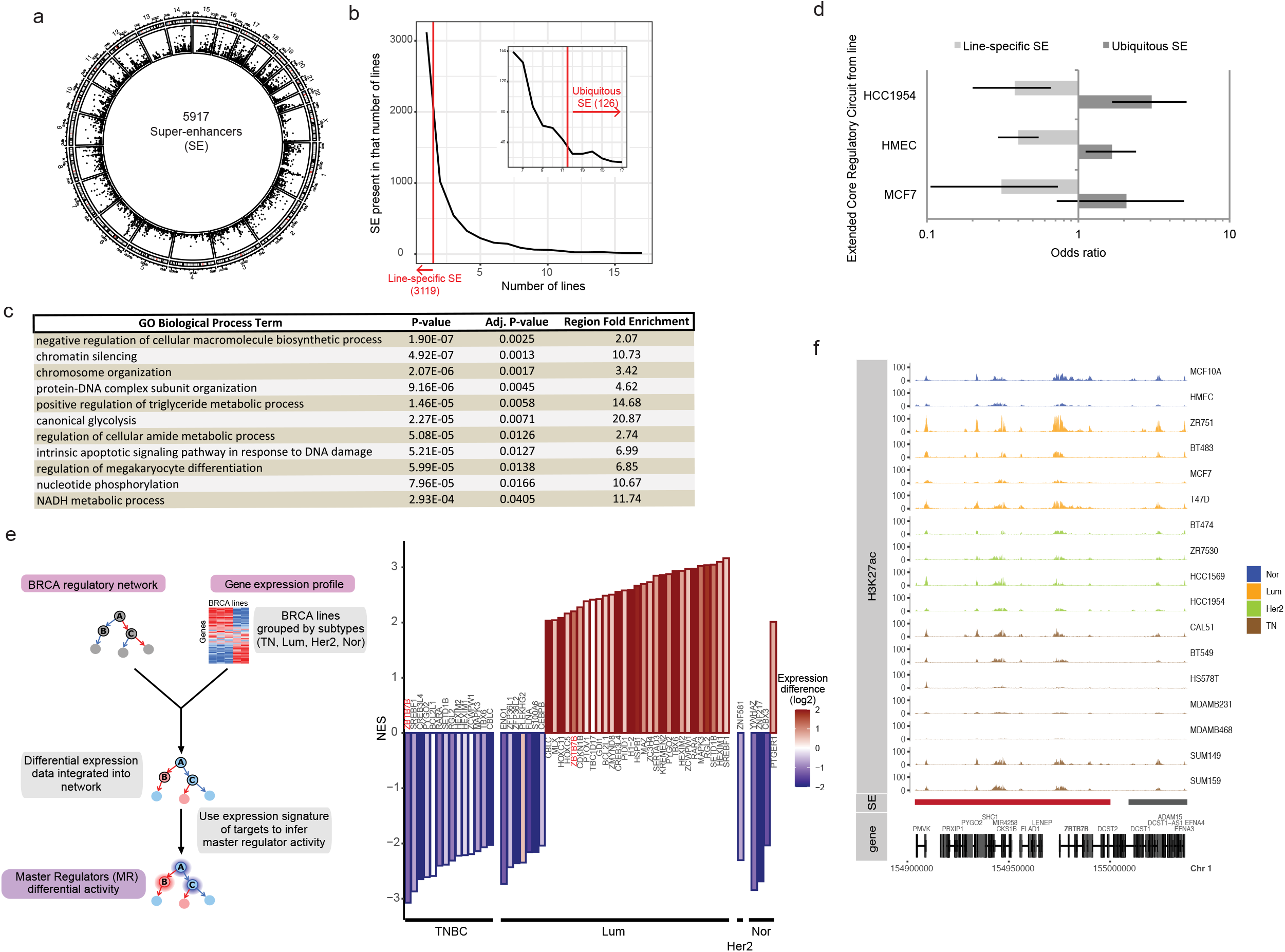
Combining ubiquitous SE identification with subtype-specific master regulator activity analysis helps identification of candidate key regulators of breast cancer cell biology. **a)** Circo plot displaying the genome distribution of the whole set of 5917 SEs that resulted from the analysis of H3K27ac density from ChIP-seq data from 17 non-transformed mammary epithelial and BCC lines. Each of the boxes in the inner circle represents an instance of a SE in one cell line. **b)** Number of SEs as a function of the number of cell lines where each SE is present. Line-specific SEs, present in only one cell line, are shown left to the red line. The inset shows a magnification of the last part of the plot (from 6 to 17 cell lines). Ubiquitous SEs are present in more than two thirds of the cell lines (12 or more), right to the red line. **c)** Representative significant GO Biological Process terms enriched for genes associated with ubiquitous SEs vs all SEs, analyzed by GREAT. The full list of terms is shown in **Suppl. Table 4. d)** Enrichment of ubiquitous SEs and depletion of line-specific SEs in the set of extended core regulatory circuitry (CRC) genes of three breast cell lines. The plot shows the odds-ratio with confidence intervals for a Fisher’s exact test (p-values: MCF7: ubiquitous SEs = 0.127, line-specific SEs = 0.0036; HCC1954: ubiquitous SEs = 0.0002, line-specific SEs = 0.0002; HMEC cells: ubiquitous SEs = 0.0089, line-specific SEs = 8.9×10^−11^). **e)** Analysis of differential activity for ubiquitous SE-associated master regulators (MR) of expression in breast cancer. Left: scheme of VIPER analysis. Subtypes: TNBC, Lum, HER2+ and normal (Nor). Right: regulators with a normalized enrichment score (NES) from VIPER analysis of more than 2 in absolute value, shown for each contrast (lines of the indicated subtype vs all the other lines). The color scale of the bars shows the change in expression of each MR for the indicated contrast. *ZBTB7B* (ThPOK) is indicated in red in the two contrasts where it appears (TNBC and Lum). **f)** Genomic context of *ZBTB7B* and its associated super-enhancer. The H3K27ac ChIP-seq signal is shown for all 17 breast cancer cell lines, grouped and colored by subtype.

While many SEs within this set are only found in one cell line (cell line-specific SE, 3119), we identified a subset (126) of ubiquitous SEs that are common across at least two-thirds of the cell lines (**Fig. 1b**). GREAT analysis revealed that genes associated with ubiquitous SEs, unlike cell line-specific SEs, are significantly enriched in terms related to gene expression regulation, chromatin remodeling, differentiation, metabolism and apoptosis (**Fig. 1c** and **Table S4**). Ubiquitous SEs are also specifically linked to core regulatory circuitry genes (**Fig. 1d**, dark grey bars) defined in Saint-André *et al* (24) as important in breast cancer cell biology. In conclusion, in contrast to cell line-specific SEs, we found that ubiquitous SEs are associated with key regulatory elements in breast cancer cell biology. Therefore, we focused on this subset for further analysis.

### ThPOK/*ZBTB7B* as a breast cancer master regulator repressed in TNBC

Using the ubiquitous SE set and the RNA-seq data from BCCs (27) to prioritize key genes and cellular states (**Table S5**), we identified key transcriptional regulators with subtype-specific activity. We employed VIPER (28) to analyze a breast cancer gene regulatory network built from The Cancer Genome Atlas (TCGA) transcriptomic data (ARACNe-BRCA) (29). This revealed the activity patterns of 4438 breast cancer master regulators (MRs) by monitoring their downstream transcriptional targets expression (i.e. regulons) (**Fig. 1e**). By comparing each subtype (Lum, HER2+, TNBC and normal (Nor)) to the rest, we calculated a normalized enrichment score (NES) for each MR in each subtype (**Table S6**). We focused on MRs with differential activity of absolute NES > 2 and genes associated with ubiquitous SEs, identifying 41 MR candidates (**Fig. 1e**). Many showed differential activity that could be due in part to variations in their own expression (**Fig. 1e**, compare activity (NES) vs RNA level). Notably, ThPOK (*ZBTB7B*), a CD4^+^ lineage commitment transcription factor, was significantly under-active in TNBC and over-active in luminal cell lines. This gene is linked to a large ubiquitous SE (**Fig. 1f**, red bar), though the H3K27ac signal near the transcriptional start site is lower in TNBC lines (**Fig. 1f, Fig. S1d-e**). To confirm SE regulation of ThPOK, we inhibited BRD4, an essential SE component (7, 17, 30-32), using JQ1 or iBET-151. Acute BRD4 inhibition (6h) reduced *ThPOK* expression (**Fig. S2a**) without affecting the housekeeping gene *ACTB* (**Fig. S2b**), or ectopic *ThPOK* expression through a stable integrated lentiviral vector (**Fig. S2c**). In addition, ThPOK fits the profile of a master regulator: it regulates gene expression, recruits chromatin repressors (33, 34), and influences cell lineage phenotypes (20, 35). Given its oncogenic potential within the POK protein family, we investigated the specific role of ThPOK in breast cancer.

### ThPOK is upregulated in luminal breast cancer cells and tumors

Quantitative RT-PCR analysis of ThPOK expression in 21 cell lines revealed subtype-associated differences in mRNA levels: highest in luminal cell lines, lowest in TNBC lines, and intermediate and more heterogeneous in HER2+ cell lines (**Fig. 2a**). This was consistent with RNA-seq data from various cell lines (**Fig. 2b**), indicating that despite the ubiquity of the SE linked to ThPOK, subtype-specific regulatory mechanisms influence its expression, resulting in quantitative mRNA differences. ThPOK protein expression, assessed by immunofluorescence, showed a similar subtype-specific pattern (**Fig. 2c**). In patient samples, RNA-seq profiles from TCGA confirmed higher *ThPOK* expression in breast tumors compared to adjacent-normal tissue (**Fig. 2d**). Luminal hormone receptor + (HR+) and HER2+ tumors exhibited higher *ThPOK* mRNA levels than TNBC tumors (**Fig. 2e**). Detailed TCGA analysis using PAM50 classification (4), and the hormone receptor status determined by immunohistochemistry (**Fig. S3a-b**) showed the highest *ThPOK* levels in PAM50 LumA and HER2+ groups (**Fig. S3a**), with lower levels in more aggressive PAM50 LumB tumors (**Fig. S3a**). ER^+^ tumors had the highest *ThPOK* levels, with HER2+ tumors showing the highest levels in the ER^-^ group (**Fig. S3b**). Several CpG sites in the 5’ UTR of *ZBTB7B* exhibited increased methylation in TNBC tumors, suggesting transcriptional repression by CpG methylation (**Figs. S3c-e)**. Supporting ThPOK’s role in breast cancer, the Human Protein Atlas showed that breast cancer ThPOK protein and mRNA levels ranked as the top and second-ranked respectively (**Figs. S4a-b**).

**Figure 2.**
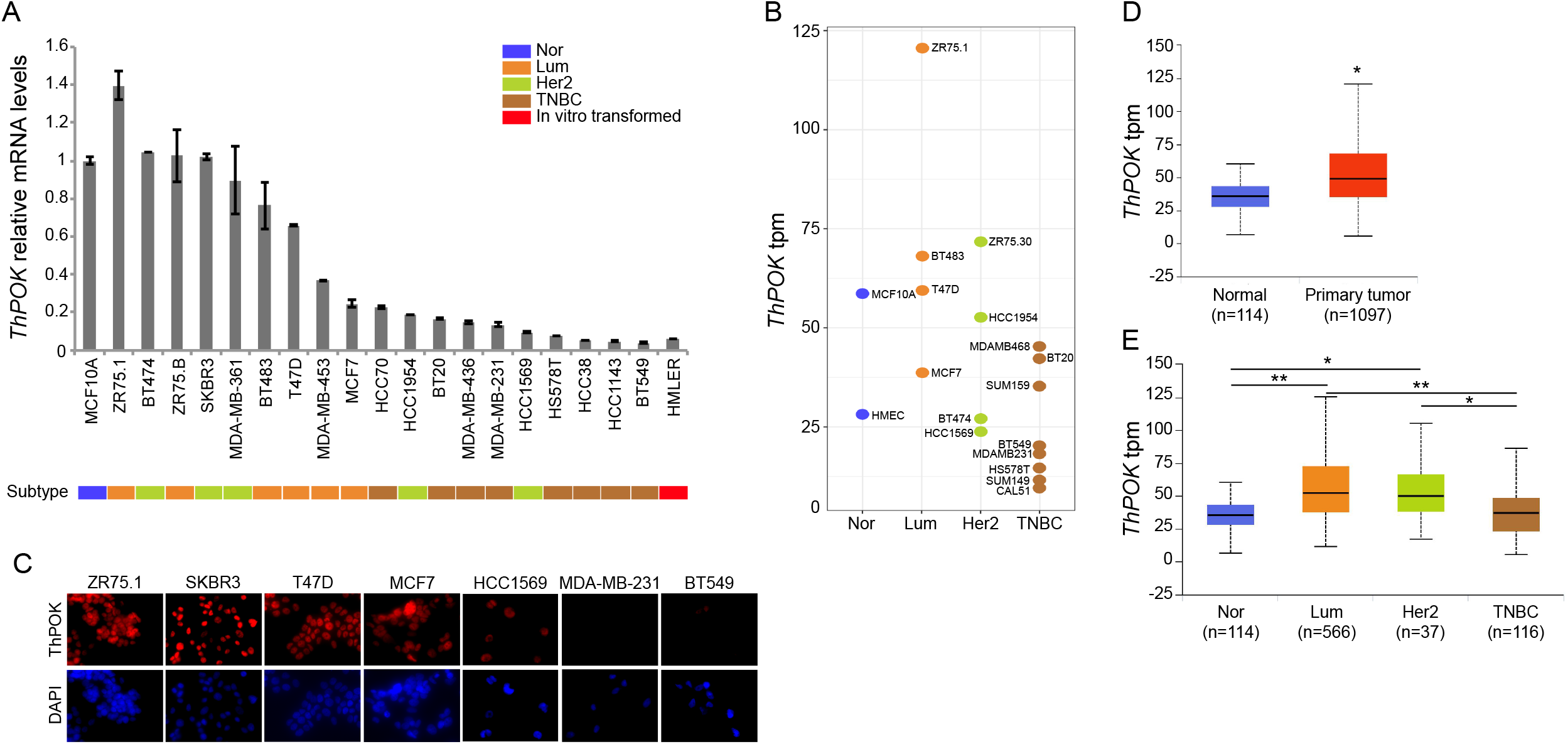
*ThPOK* expression is highest in breast cancer cell lines and tumors of the luminal subtype. **a)** *ThPOK* expression in various BCC lines measured by qRT-PCR. Error bars: SD of ≥3 biological replicates. All values are referred to the average expression level in MCF10A cells. Breast cancer subtypes are colored as in Fig. 1: blue, Nor; orange, Lum; green, HER2+; brown, TNBC; red, *in vitro* transformed cells. **b)** *ThPOK* expression extracted from RNA-seq datasets listed in **Suppl. Table 6. c)** ThPOK expression detected by immunofluorescence (red channel). Nuclei were counterstained with DAPI. Images are shown at 40X magnification. **d) & e)** *ThPOK* expression in primary tumors (TCGA dataset). **d)** *ThPOK* expression is higher in primary tumors than in normal tissue, p=1.62×10^−12^. **e)** *ThPOK* expression in breast cancer subtypes. Luminal tumors express the highest *ThPOK* levels compared to normal tissue (p<10^−12^), followed by HER2+ tumors (p=2.30×10^−5^). *ThPOK* expression in TNBC tumors is not significantly different to the levels in normal tissues. In **d & e** in parenthesis: number of tumors/normal tissues included in each subtype. These plots were generated using the UALCAN portal (http://ualcan.path.uab.edu).

### ThPOK silencing de-represses a pro-migratory gene expression program

Although ThPOK is mainly known as a transcriptional repressor, depending on the context it can also act as an activator (33, 36-38). To understand ThPOK’s specific action, we used shRNA-mediated knockdown (kd) to reduce its mRNA levels (**Fig. 3a, S5a-d**) in luminal (T47D, MCF7 and ZR75.1) and HER2+ (BT474) cell lines with high to intermediate *ThPOK* mRNA levels (**Fig. 2a**). We initially assessed the expression levels of known ThPOK target genes in fibroblasts and T cells, which may have a role in breast cancer. ThPOK deficiency led to increased ECM gene expression (*FN1, COL1A1, COL1A2*) (**Fig. 3b**), and caused EMT-like morphological changes in most cell lines (**Fig. 3c, S5e**). Since ThPOK has features of a master regulator, we assessed the impact of its silencing on other gene sets representing different cancer hallmarks. ThPOK kd led to an increase in the expression of EMT factors (*SNAI1, SNAI2, ZEB1, ZEB2, TWIST*), basal markers and WNT/β-catenin target genes implicated in breast cancer cell proliferation (39, 40) (**Fig. 3d-f**). The protein membrane and stemness marker CD44 was also upregulated as shown by qRT-PCR and flow cytometry analysis (**Fig. 3g, S5c**). Consistent with this, using TCGA breast tumor’s genomic data, we found that low levels of ThPOK correlate with a higher RNA-based stemness score (41) (**Fig. S5d**). Simultaneously, we observed a mild decrease in the expression of epithelial markers, especially CDH1, and deregulation of *ESR1* (**Fig. 3d, h & i**). Consistent with a general alteration of their transcriptional programs, ThPOK-deficient luminal cells exhibited an enhanced migratory phenotype (**Fig. 3j-k** and **S5e**).

**Figure 3.**
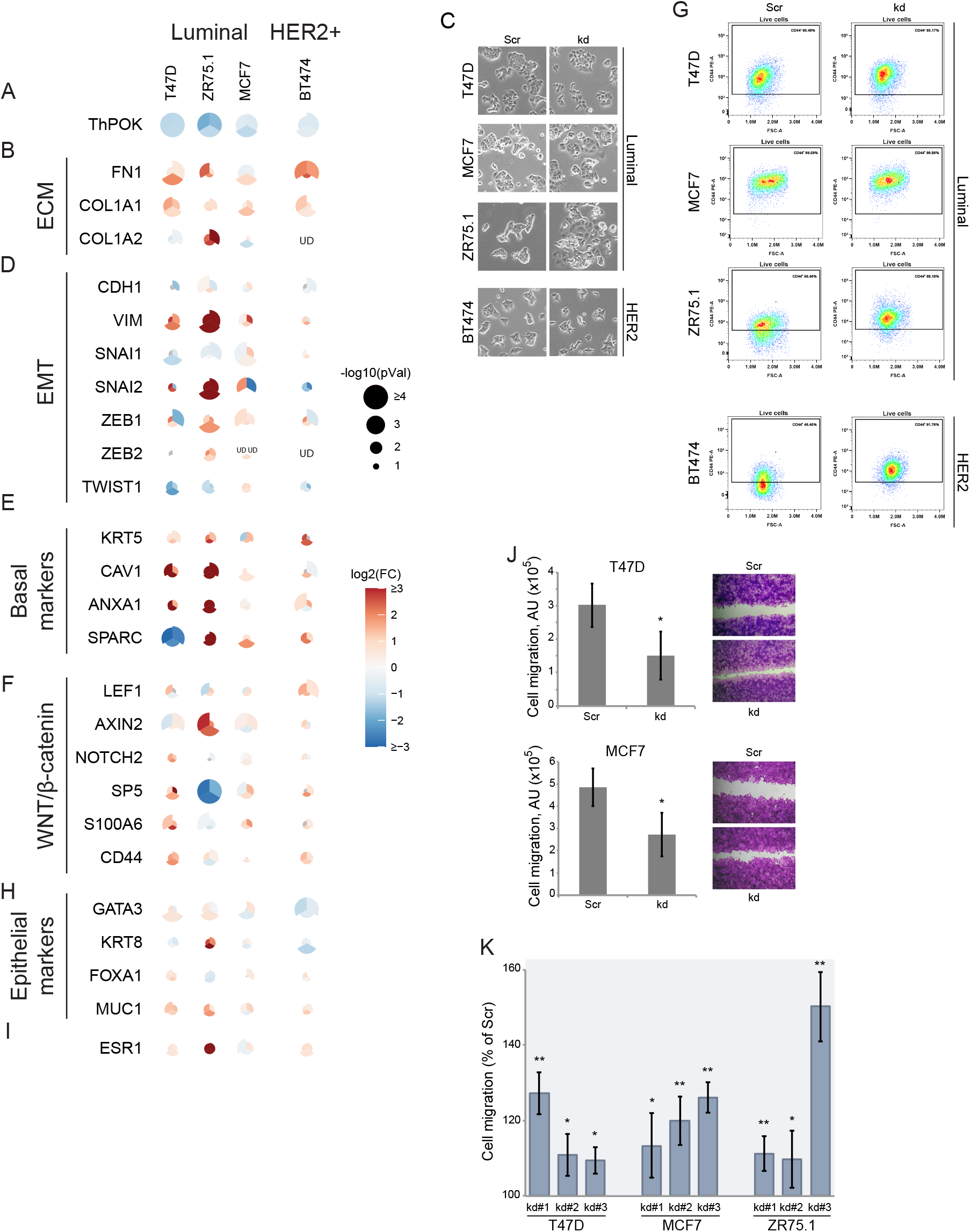
*ThPOK* knockdown increases expression of ECM, EMT and WNT/β-catenin genes in luminal and HER2+ BCCs. **a, b, d-f, h-i)** Gene expression levels measured by qRT-PCR. Since three different shRNA against *ThPOK* were used, the circle for each gene is divided in thirds, one for each shRNA. The color of each third indicates the log_2_(fold-change) and the size the -log_10_(p-value), for a *t*-test relative to the control shRNA (shScr). Each gene was assessed in ≥3 biological replicates. Differential expression is shown in color scale as a log2 of the fold change, and p-values by the size of the circle as –log10. **c)** Morphological changes associated with ThPOK deficiency in Luminal and HER2+ cells (10X) magnification. Scale: each side of the square is 400µm. **g)** Flow cytometry analysis showing CD44 expression in Luminal and HER2+ scramble (Scr) and ThPOK-kd cells. **j-k)** Migration assays. **j)** Scratch assay showing increased migratory features of ThPOK-kd cells compared with control (scr) cells. Error bars represent the SD of ≥3 biological replicates. T47D cells p=4.07×10^−19^, MCF7 cells p=8.087×10^−8^. **k)** Boyden chamber assay. Error bars represent the SD of ≥3 biological replicates, p *<0.05, **<10^−4^.

### ThPOK expression induces a “luminal-like” gene signature in HER2+ and TNBC cell lines

Based on our previous results, we hypothesized that overexpressing ThPOK in HER2+ and TNBC cells, which have lower endogenous ThPOK levels, would cause opposite changes in gene expression patterns and cellular phenotype. We transduced HCC1954 (HER2+), and HCC1143, BT549 and MDA-MB-231 (TNBCs) with a lentivirus encoding ThPOK variant #5, the variant expressed in BCCs (**Fig. S6a-b)**. ThPOK overexpression (ThPOK^OE^) was confirmed by qRT-PCR and Western Blot (**Fig. 4a & Fig. S7a**). As predicted, ThPOK overexpression decreased ECM genes expression and increased epithelial markers including *CDH1*, while decreasing the expression of genes associated with migration and invasiveness, such as EMT markers (*VIM, SNAI1, SNAI2, ZEB1, ZEB2* and *TWIST*), WNT/β-catenin target genes, the stemness marker *CD44* and *ESR1* (**Figs. 4b-g**). These changes are accompanied by a mesenchymal to epithelial transition-like phenotype and reduced migratory features (**Figs. 4h-i & S7b**).

**Figure 4.**
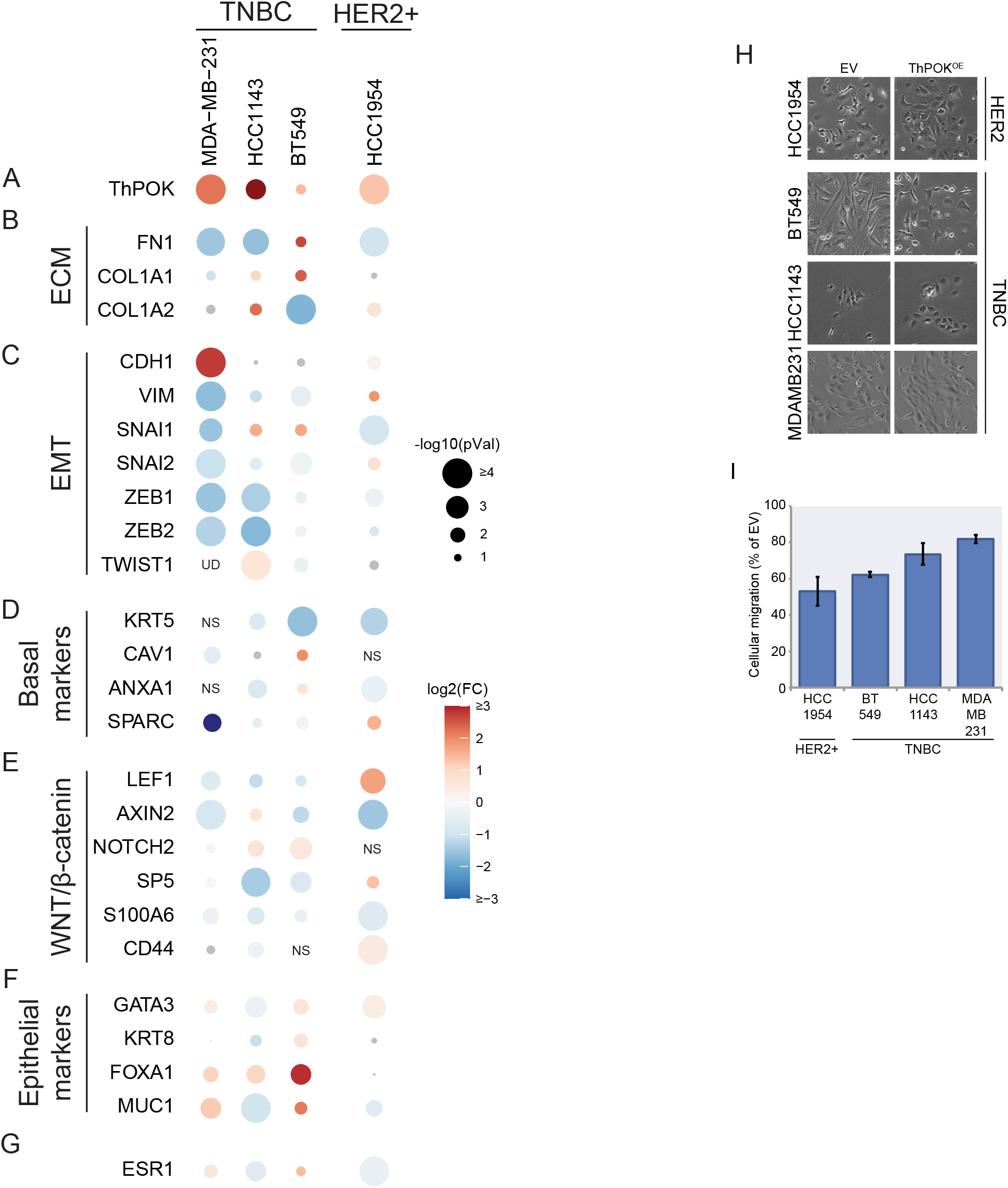
*ThPOK* overexpression represses ECM, EMT and WNT/β-catenin genes in HER2+ and TNBC BCCs. **a-g)** Gene expression levels measured by qRT-PCR. The log_2_(fold-change) in expression of each gene is shown in color scale and p-values by the size of the circle as –log_10_, relative to the empty vector control cells (EV). Each gene was assessed in ≥3 biological replicates. **h)** Morphological changes associated with ThPOK overexpression in HER2+ and TNBC cells (10X) magnification. Scale: each side of the square is 400µm. **g)** Boyden chamber migration assays depicted as percentage of that of EV cells, error bars represent the SD of ≥3 biological replicates, *p<10^−6^.

Taken together, these data indicate that ThPOK functions as a transcriptional repressor of genes that are critical for breast cancer phenotypes associated with invasiveness and aggressiveness.

### ThPOK inhibits the TGFβ pathway but independently regulates ECM genes

Since ThPOK inhibits the expression of *FN1* and *COL1A1*, which are positively regulated by TGFβ signaling (42), we hypothesized that ThPOK could be negatively regulating the TGFβ pathway, a central regulator of ECM remodeling and fibrosis. Indeed, when we analyzed the levels of TGFβ/SMAD target genes in both, breast cancer cell lines from the Cancer Cell Line Encyclopedia and breast tumors from the TCGA, we saw an inverse correlation between the expression of ThPOK and of TGFβ target genes (**Fig. 5a**). By measuring TGFβ (*TGFB1*) mRNA levels in ThPOK-deficient (**Fig. 5b**) and ThPOK^OE^ cells (**Fig. 5c**) by qRT-PCR, we confirmed that ThPOK is actually a repressor of TGFβ mRNA expression in BCCs. We hypothesized that if ThPOK represses ECM gene expression directly and not through the repression of TGFβ, high levels of ThPOK should counteract the induction of ECM genes by TGFβ treatment. Consistent with this notion, while treatment of luminal cancer cell lines with recombinant TGFβ (rTGFβ) increased *FN1* and *COL1A1* expression, the magnitude of the effect was smaller, and a longer treatment was needed in T47D cells that have high endogenous ThPOK levels compared to MCF7 cells with lower levels (**Figs. 5d** and **S8a**). Similarly, ectopic ThPOK overexpression in TNBC counteracted the inducing effects of rTGFβ treatment compared to empty vector (EV) control cells for the assessed ECM genes (**Figs. 5e** and **S8b**). These results suggest that in both scenarios, high levels of endogenous and/or ectopic ThPOK can partially counteract the inducing effects of TGFβ signaling in the expression of ECM genes, suggesting a direct regulatory role.

**Figure 5.**
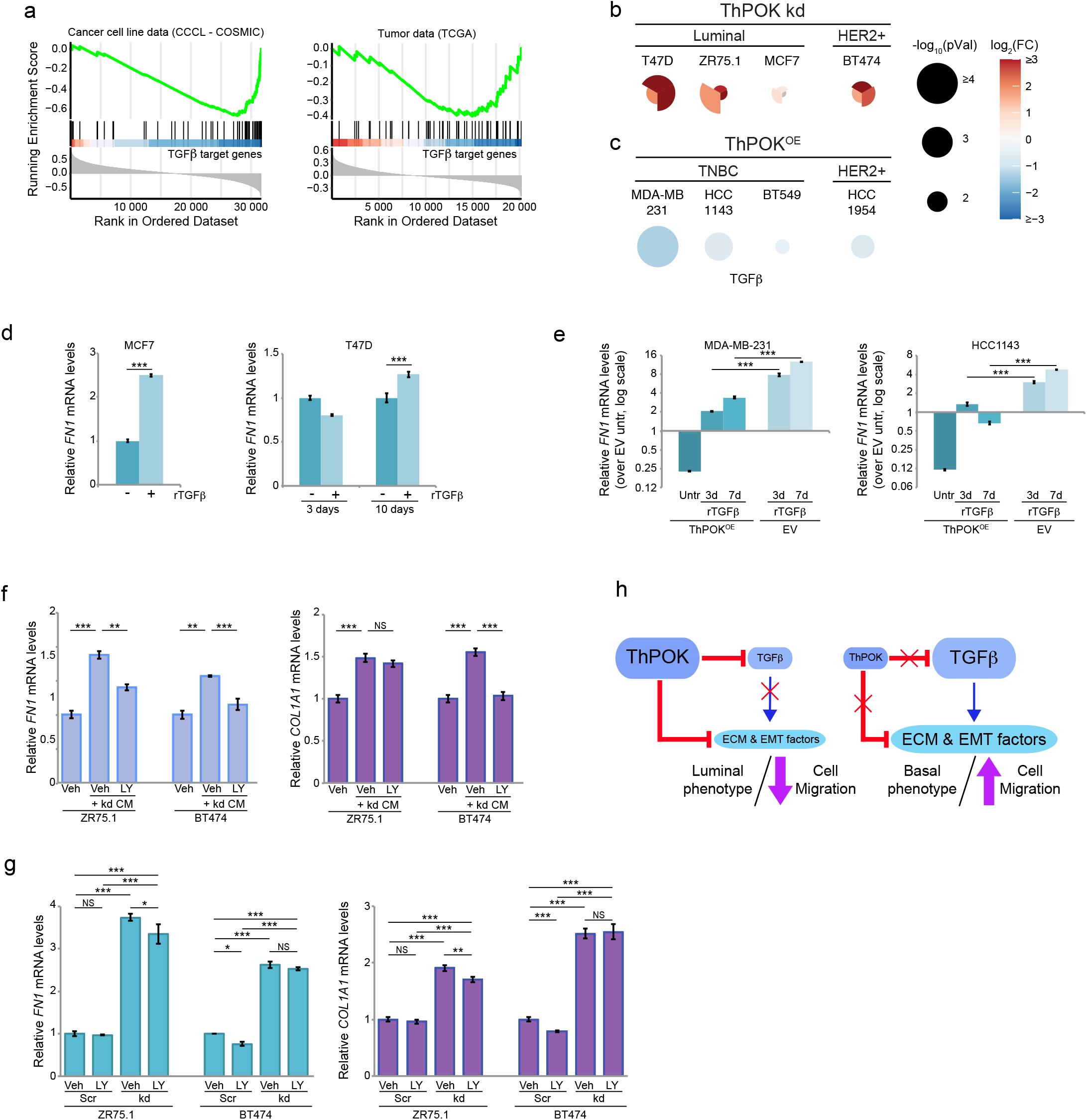
ThPOK represses TGFβ expression in breast cancer cell lines. **a)** GSEA analysis of the relationship between ThPOK and TGFβ target genes expression levels using transcriptomic data from CCCL (breast cancer cell lines) or TCGA (breast tumors). **b) & c)** TGFβ (*TGFB1*) mRNA levels measured by qRT-PCR. **b)** TGFβ levels in ThPOK-deficient luminal and HER2+ BCCs. *TGFB1* expression in each independent shRNA-containing cells is depicted as a triangle and the fold change is relative to that of the control shRNA (shScr). Each gene was assessed in ≥3 biological replicates. Differential expression is shown in color scale as a log_2_ of the fold change, and p-values by the size of the circle as –log_10_. **c)** TGFβ levels in HER2+ and TNBC cells overexpressing ThPOK (ThPOK^OE^). *TGFB1* expression is shown as the fold change relative to that of the empty vector control cells (EV). Each gene was assessed in ≥3 biological replicates. Differential expression is shown in color scale as a log_2_ of the fold change, and p-values by the size of the circle as –log_10_. **d)** *FN1* relative mRNA levels measured by qRT-PCR in MCF7 cells after 3 days treatment with recombinant human TGFβ (rTGFβ), and T47D cells after 3 or 10 days treatment with rTGFβ. **e)** *FN1* relative mRNA levels measured by qRT-PCR in MDA-MB-231 and HCC1143 cells with ThPOK^OE^ or EV, untreated (Untr) or treated with rTGFβ for 3 or 7 days. **f)** *FN1* and *COL1A1* relative mRNA levels measured by qRT-PCR in ZR75.1 and BT474 cells treated with DMSO (Veh), Veh + conditioned media (CM) from ThPOK kd cultures, and CM + TGBRI inhibitor (LY2157299, LY). **g)** *FN1* and *COL1A1* relative mRNA levels measured by qRT-PCR in Scr and ThPOK kd (kd) ZR75.1 and BT474 cells, treated with DMSO (Veh) or TGBRI inhibitor (LY). For **d**-**g**, all expression values are depicted relative to the mean of the corresponding controls. Error bars represent the SD of ≥3 biological replicates. NS: non-significant; *: *p* < 0.05; **: *p* < 0.005; ***: *p* < 0.0005; 2-samples *t*-test. **h)** Proposed model showing the direct and indirect regulation of extracellular matrix, EMT and TGFβ/SMAD target genes’ expression by ThPOK. The size of the balloons represents the relative contribution (not in scale) of each factor to the cellular phenotypes.

TGFβ is produced as a latent complex and is activated extracellularly by many factors (42). To test whether ThPOK-deficient cells are able to activate the produced TGFβ, control cells were exposed to conditioned media collected from ThPOK kd cultures in the presence or absence of a TGFβ receptor I (TβRI) inhibitor (LY2157299), and expression of *FN1* and *COL1A1* genes was assessed. Conditioned media from ThPOK kd cells significantly increased ECM gene mRNA levels in luminal/HER2+ cells, and this effect was mostly mitigated by TβRI inhibition (**Fig. 5f**), indicating that ThPOK-deficient cells produce and are able to activate TFGβ. Although we have shown above that ThPOK counteracts the inductive effects of TGFβ signaling in the expression of ECM genes, these results could also indicate that the produced TGFβ may influence ECM gene expression in an autocrine manner rather than being solely driven by the lack of ThPOK. To assess this, we inhibited TβRI in ThPOK kd cells and evaluated *FN1* and *COL1A1* expression. Although TβRI inhibition slightly reduced the expression of TGFβ target genes, it did not fully revert their levels to those of the original scramble control cells (**Fig. 5g**, compare Veh vs kd+LY). This implies that while TGFβ produced by ThPOK kd cells may autocrinally contribute to some extent to the increase in ECM gene expression, the primary driver is the absence of ThPOK’s repressive functions. In summary, ThPOK is a key transcriptional regulator that limits breast cancer cells plasticity by repressing TGFβ□□ECM genes, and stemness and mesenchymal factors, thereby promoting an epithelial, non-migratory phenotype (**Fig. 5h**).

## DISCUSSION

Epigenetic dysregulation is a hallmark of cancer, and the identification of SEs has aided in the discovery of unexpected dependencies and novel drug targets (8, 32). In this study we employed a multi-omic approach that integrates SE profiling and transcriptomic analysis of 17 normal mammary and breast cancer cell lines with a breast cancer-specific gene regulatory network, revealing ThPOK/*ZBTB7B* as a subtype-specific master regulator in breast cancer. The *ZBTB7B* gene was among the few breast cancer transcription factor genes associated with a ubiquitous SE despite displaying varying expression levels. This may indicate that SEs impart complex regulatory architecture rather than solely controlling gene expression, potentially allowing chromatin accessibility at the ThPOK locus permitting fine-tuning to adapt to the changing demands of the lactating mammary gland. Our discovery of ThPOK as a master regulator is further supported by its role as a methylation modulator gene in breast invasive carcinoma (43). An integrated analysis of DNA-methylation and RNA-seq data from ∼500 breast invasive carcinoma samples highlights ThPOK as a hub node within a gene module containing several genes regulated by WNT/β-catenin and TGFβ (43).

ThPOK’s physical interaction with histone deacetylases (HDACs), and non-transcribed genes suggests that ThPOK primarily functions as a transcriptional repressor (33, 36). Consistent with these findings, ThPOK represses genes involved in ECM composition, as well as transcription factors implicated in the EMT and WNT/β-catenin signaling pathways. These effects coincided with cellular morphological and behavioral changes indicative of enhanced migratory capacity in breast cancer cells. In addition, we identified a common set of genes regulated by ThPOK and polycomb repressor complex 2 (PRC2) in transformed mammary epithelial cells (44). Dysfunction of PRC2 and ThPOK deficiency, both lead to a quasi-mesenchymal state, characterized by de-repression of many EMT and basal genes and retention of some epithelial features. We are currently investigating whether ThPOK associates with polycomb repressor family of proteins to exert its repressive functions.

The ECM influences cancer cell behavior through chemical and mechanical signals, with changes in collagen composition linked to tumor invasion and aggressiveness (45). Recent studies have shown that collagen deposition and organization regulate dormancy in disseminated breast cancer cells and immune exclusion in TNBC (46, 47). Given ThPOK’s repression of ECM genes, the regulation of ThPOK expression could profoundly impact the ECM and consequently alter cancer cell behavior. This is also consistent with the fact that ThPOK regulates *TGFB1* mRNA and TGFβ signaling pathways play a critical role in tumor formation, progression and metastasis (40). Notably, *FN1* and *COL1*, all regulated by TGFβ (48, 49), exhibit coordinated expression in breast cancer lymph node metastases (49). These proteins form networks that promote a spindle-like phenotype, significantly enhancing tumor cell migration. The observation that ThPOK also regulates these same genes raises the possibility that this regulation occurs indirectly through the modulation of TGFβ. However, our findings and previous studies suggest a direct and independent role of ThPOK in the regulation of ECM and *TGFBI* genes (18, 19, 50).

A complex interplay among GATA3, RUNX and ThPOK determines T cell fate, with GATA3 as an upstream positive regulator of ThPOK in the thymus (51) and ThPOK influencing GATA3 expression in breast cancer cells. GATA3, a critical regulator of luminal differentiation during mammary gland development, also acts as a key transcription factor in maintaining an epithelial phenotype in luminal breast cancers (52-54). Interestingly, ThPOK levels are similarly elevated in this subtype, supporting the idea of a functional interaction between GATA3 and ThPOK where they may collaborate to maintain luminal identity in both normal and cancerous cells (52, 53). Moreover, GATA3 participates in a positive feedback loop with ERα in breast cancer (53), and ThPOK interacts with ERα in the nuclei of breast cancer cells (55). Exploring the interplay among ThPOK, GATA3 and ERα could provide insights into breast cancer progression, and luminal differentiation and maintenance.

A subset of ductal carcinoma in situ (DCIS) cases progress to invasive ductal carcinoma (IDC), posing clinical challenges in identifying patients requiring a more aggressive treatment. Investigating ThPOK repression as a mechanism facilitating the acquisition of more aggressive phenotypes could shed light on the transition to invasive disease.

In conclusion, our multi-omic approach uncovered ThPOK, known as a CD4^+^ lineage commitment factor, as a breast cancer master regulator, with manipulation of ThPOK levels revealing a network of target genes enriched in cellular plasticity programs. Understanding how cancer cells regulate repressive transcription factors like ThPOK, which absence can become a tumor-promoting factor, is critical for mitigating carcinogenesis and tumor progression.

## MATERIALS AND METHODS

### Cell lines, culture conditions, constructs and lentivirus production and transduction

HCC1954, BT20, BT483, HCC1569, and BT474 breast cancer cell lines were bought from ATCC. MDA-MB-361 and MDA-MB-453 were bought from UC Berkeley Cell Culture laboratory. ZR75.1, MDA-MB-436, and ZR75.B were kindly given by Dr. Desprez. MCF10A, MCF7, T47D, MDA-MB-231, and SKBR3 were kindly given by Dr. Kohwi-Shigematsu. HMLER cells were kindly provided by Dr. Weinberg. All cell lines tested negative for Mycoplasm, were grown at 5%CO_2_ with all media was supplemented with 10% FBS and 1X Penicillin/Streptomycin. BT20, BT474, HS578T, MCF7, MDA-MB-231, MDA-MB-453, MDA-MB-361, MDA-MB-436, and SKBR3 were grown in DMEM; BT483, BT549, HCC1143, HCC38, HCC70, HCC1569, HCC1954 and ZR75.1, MDA-MB-436, ZR75.B, and T47D in RPMI. MCF10A and HMLER cells were grown as described (56, 57).

### ThPOK construct

*ThPOK* mRNA variant % (NCBI Reference Sequence: NM_001256455.2) was amplified by PCR from cDNA and cloned into pENTR1A-DS vector (Invitrogen) using SalI and NotI sites, then sequenced for accuracy. The pENTR1A-ThPOK vector was recombined into pLenti-CMV-DEST-hygro (W117-1) (provided by Dr. Campeau (58)) using Clonase II (Thermofisher, cat# 11791).

### Lentivirus production and transduction

shRNA-expressing lentiviral vectors for ThPOK knockdown were purchased from Open Biosystems/Dharmacon (RHS4533-EG51043). Lentiviruses containing shRNA or ThPOK constructs were produced as described in Rodier *et al* (59). Transductions were done overnight with 4µg/ml of Polybrene™. 48h later cells were selected with 2µg/ml puromycin or 400µg/ml hygromycin.

### Chromatin immunoprecipitation and next generation sequencing (NGS)

Chromatin preparation and immunoprecipitation were performed as described in (60) using a H3K27ac antibody (Abcam cat# Ab4729). ChIP-seq libraries were prepared using Illumina protocols, quantified with Kapa Biosystems kits, pooled and sequenced on a HiSeq4000 as 50-base single reads.

### NGS data analysis

RNA-seq data were quality controlled with FastQC. Reads were aligned and transcript counts were obtained using kallisto with 100 bootstraps (61). Master regulator activity inference from RNA-seq data was performed with VIPER (28). ChIP-Seq data were quality controlled with FastQC, mapped using Bowtie 2.0, and H3K27ac peaks and SE were obtained using MACS2 and ROSE2 algorithms (7, 8, 25). For more detail refer to supplementary methods.

### Immunofluorescence

ThPOK immunofluorescence was performed as previously described (62), using an antibody from Novus Biologicals cat# NBP1-88077. Secondary antibody was anti-rabbit IgG ReadyProbes™ Alexa Fluor 594. Nuclei were counterstained with DAPI. Images were taken using the same exposure time using a Keyence fluorescence microscope BZ-X800 at 400X magnification.

### Western-blot

Cells were lysed and proteins separated by SDS-PAGE. Primary antibodies: ThPOK (Novus Biologicals cat# NBP1-88077); GAPDH (Cell Signaling cat# 97166); β-Actin (Abcam cat# ab6276). Secondary antibodies: anti-mouse and anti-rabbit HRP-conjugated (BioRad cat# 1705047EDU and 1662408EDU).

### RNA extraction, cDNA synthesis and qRT-PCR

RNA was purified with TRIzol followed by DNAse RNase-free digestion (Qiagen). cDNA synthesis was performed using the High Capacity cDNA Reverse Transcription kit (ABI). qRT-PCR primers sequences will be provided upon request. qRT-PCR was performed using SYBR-Green Power master mix (Thermofisher), analyzed with the QuantStudio5 software, and relative mRNA levels were calculated using the 2^^(-ΔΔCt)^ method using *GAPDH* and *HPRT1* as controls. We used a *t*-test for two-samples with equal variance (with a two-tailed distribution) to test the significance of gene expression differences.

### Flow cytometry

For CD44 staining, cells were collected, washed in FACS buffer (2% FBS in PBS), and kept at 4ºC for the rest of the protocol. Cells were stained for 1 hour with an anti-CD44-PE antibody (Cell Signaling, cat# 88151) at a 1:300 dilution in FACS buffer. After washing, cells were stained for 20 minutes with LIVE/DEAD Fixable Violet Dead (Life Technologies) at a 1:7500 dilution in PBS. After washing, CD44+ events were quantified using a Northern Light cytometer (Cytek). Data were analyzed on SpectroFlo software (Cytek).

### Scratch and Boyden chamber assays

For the scratch assay, once cells reached confluency three parallel scratches were made per well. After 48h, cells were fixed with 4% (V/V) paraformaldehyde, stained with 0.1% Crystal Violet for 20min, rinsed and air-dried. The scratches were imaged using a Nikon Eclipse TS100 light microscope under 4x objective. Multiple and overlapping images were taken along each scratch. Image J was used to measure the area unoccupied by cells, which was reported under the measurement parameter “Area”. We created Macro Plug-in with a code to analyze the images in batches. Data from individual images were grouped together per scratch. Images with highly irregular shapes and of the head and the tail region of the scratches were excluded. The assay results of each scratch were calculated by taking the mean of those grouped images, and then plotting based on different treatments. Data were obtained from at least 3 biological replicates. Boyden chamber assay was performed as described in (62) with no Matrigel and 10% FBS-media as chemoattractant. Cells on the outside bottom of the wells were fixed with 4% paraformaldehyde, stained with 0.1% crystal violet, extracted with 10% acetic acid and quantified at 590nm. Experiments were done three times in triplicates.

### Treatment with TGFβ and inhibition of TGBRI

Cells were treated with 10ng/ml of rTGFβ (Life Technologies, cat#: PHG9204) for the indicated times. To assess the activity of TGFβ produced by ThPOK kd cells, the cells were grown in RPMI/DMEM with 10% of Serum Replacement Medium (KO SRM Gibco cat# 10828-028) for 72h before the supernatant was collected, centrifuged and filtered to constitute the conditioned media (CM). shScr cells were grown in the presence of conditioned media with or without the TGBRI inhibitor LY2157299, at a 2nM concentration for 24h. To assess TGFβ’s autocrine function, cells were grown in RPMI/DMEM with 10% SRM for 24h with or without the TGBRI inhibitor LY2157299 at a 2nM concentration.

### GSEA and stemness score analyses

CCLE and TCGA datasets were used to calculate Spearman correlations between *ZBTB7B* and the rest of the genes, and the correlation-ranked genes were analyzed for enrichment of TGFβ target genes (63) using GSEA (64) within the clusterProfiler R package (65). TCGA BRCA data on ZBTB7B RNA expression and RNA-based stemness score was downloaded from Xenabrowser (66). Samples were divided in two groups based on the median value of ZBTB7B RNA expression. t-test with Welch’s correction was performed.

## Acknowledgments

We thank Diane Heditsian, our breast cancer patient advocate for her priceless support and discussions of patients’ needs to ensure we fairly represent breast cancer patients in our research, and Martín García Solá for his support and advice.

This work was supported in part by National Institutes of Health (NIH/NCI) [5K22CA163969-02], Pink Pumpkin Patch, Swim Across America and Breast Research Development Program (UCSF) grants to DPM, grants from the Agencia Nacional de Promoción de Ciencia y Tecnología of Argentina and the Universidad de Buenos Aires to IES, and by NIH/NCI Cancer Metabolism Training Program Postdoctoral Fellowship [T32CA221709] and Mentored Research Scientist Development Award to Promote Diversity [K01CA279498] to RWS.

