## Supplementary figures and images for "Super-enhancer profiling reveals ThPOK/ZBTB7B, a CD4^+^ cell lineage commitment factor, as a master regulator that restricts breast cancer cells to a luminal non-migratory phenotype"

### Supplementary Figure 1

#### Supplementary Figure 1

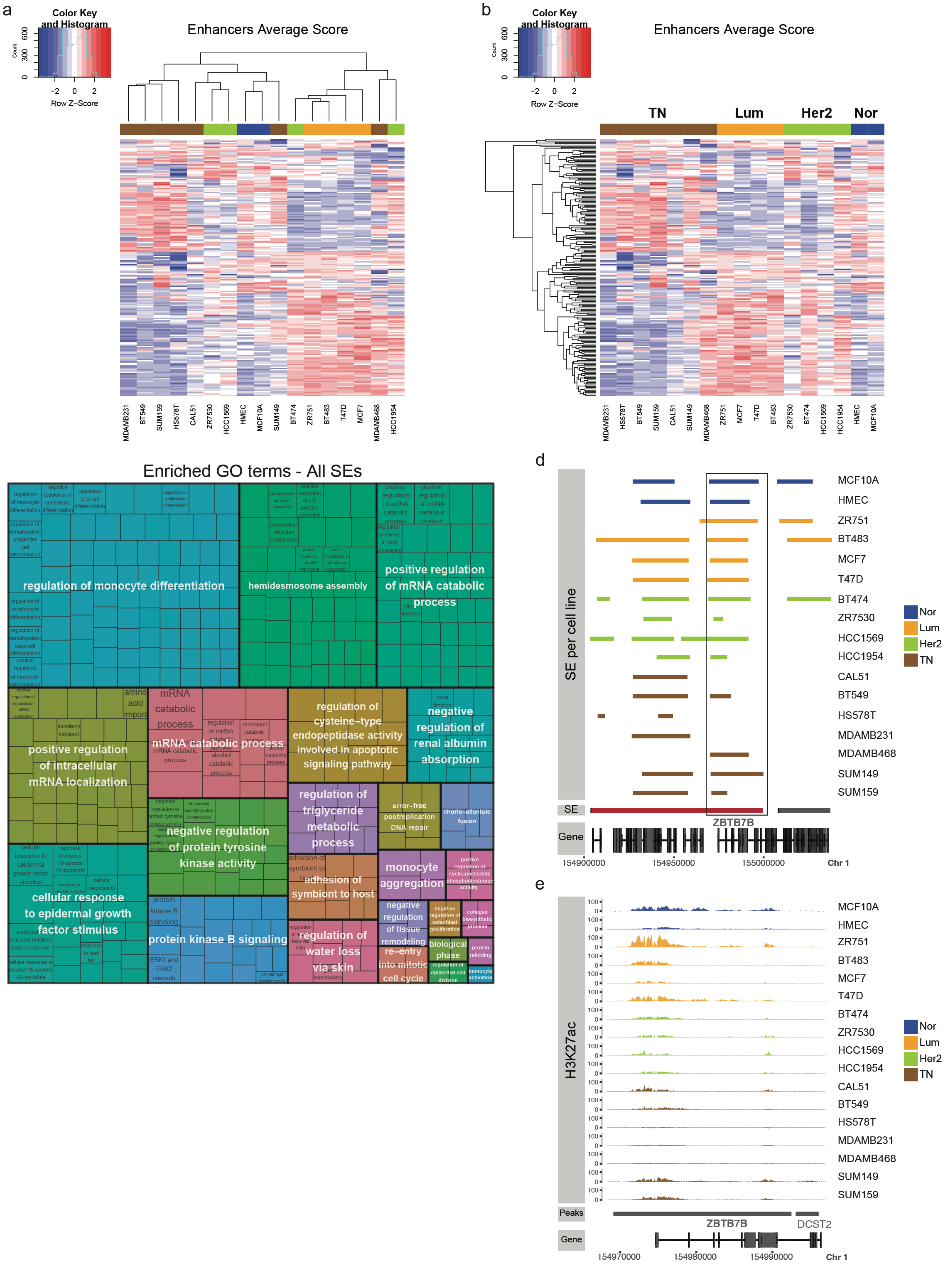

### Supplementary Figure 2

#### Supplementary Figure 2

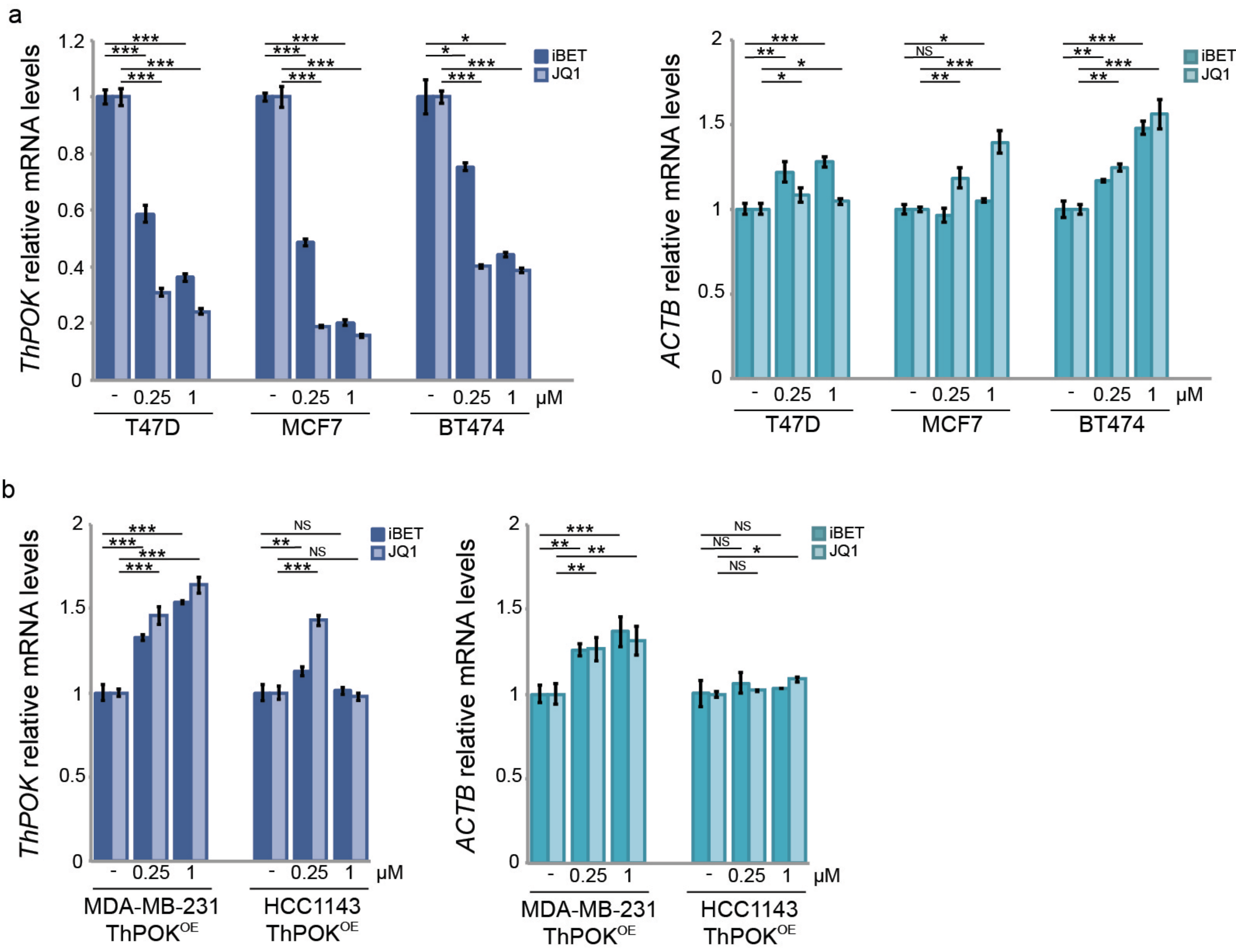

### Supplementary Figure 3

#### Supplementary Figure 3

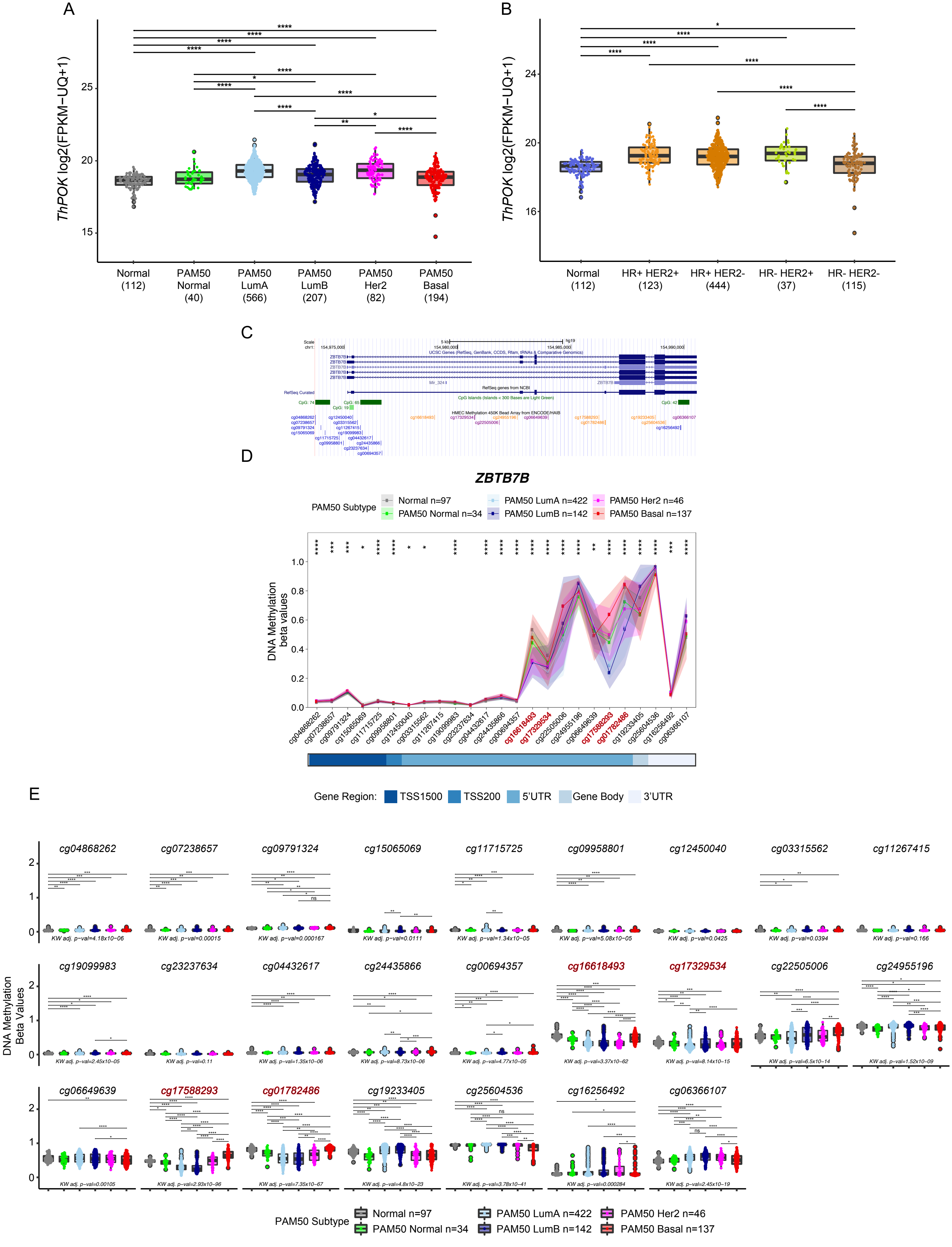

### Supplementary Figure 4

Supplementary Figure 4

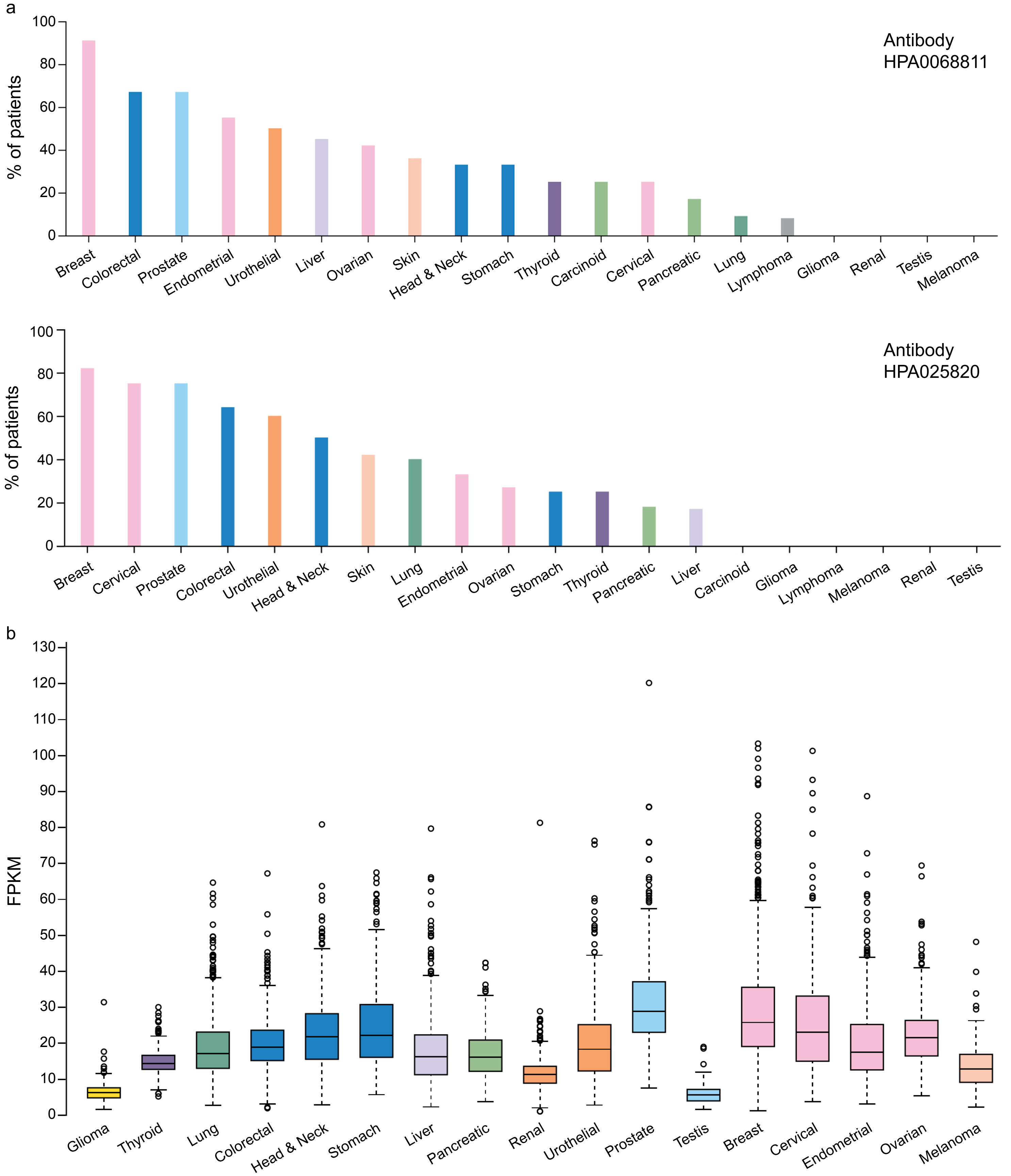

### Supplementary Figure 5

#### Supplementary Figure 5

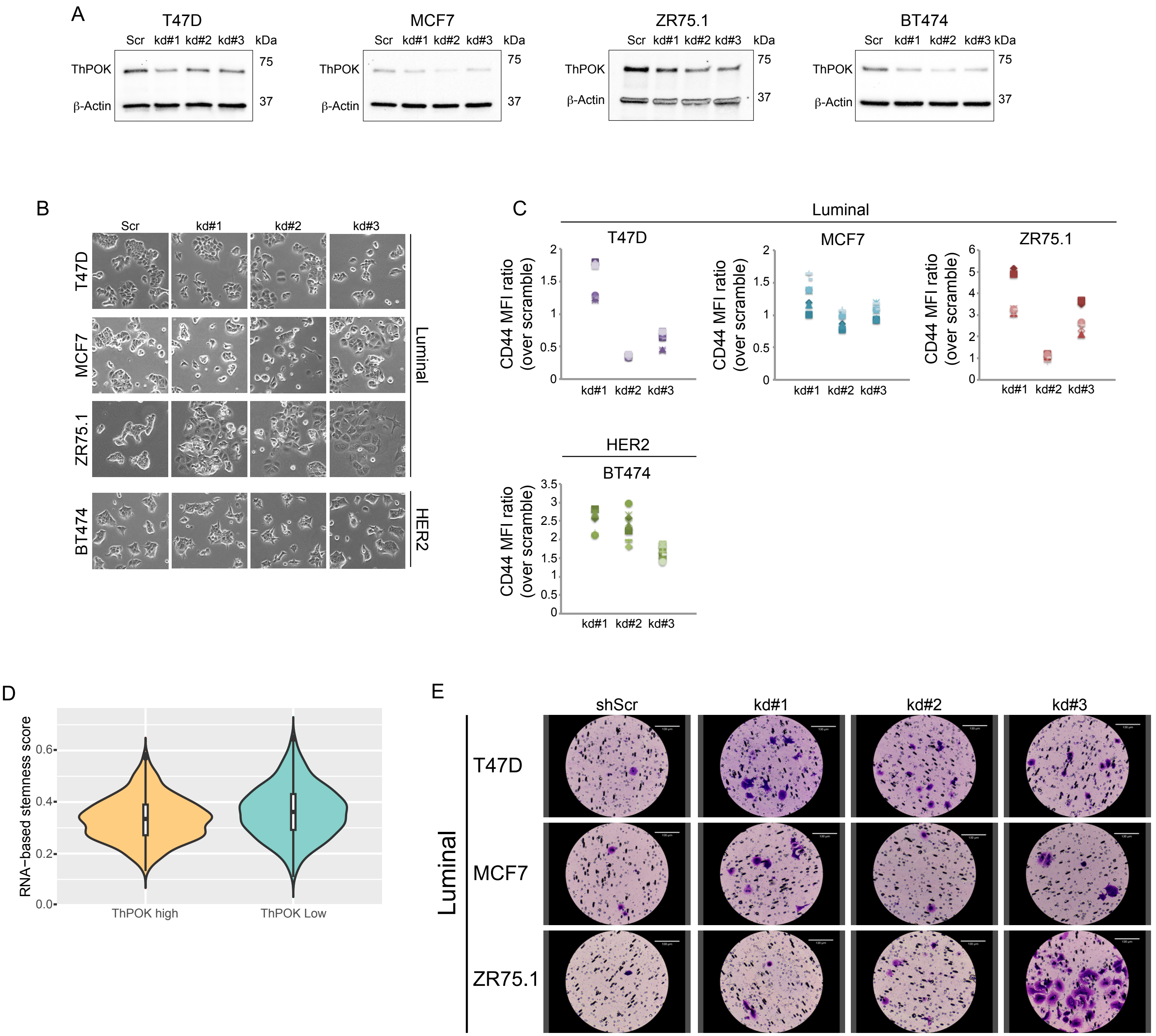

### Supplementary Figure 6

#### Supplementary Figure 6

a

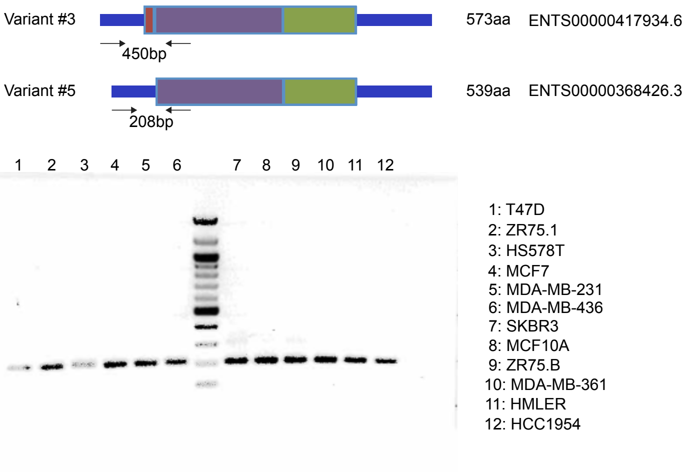


b

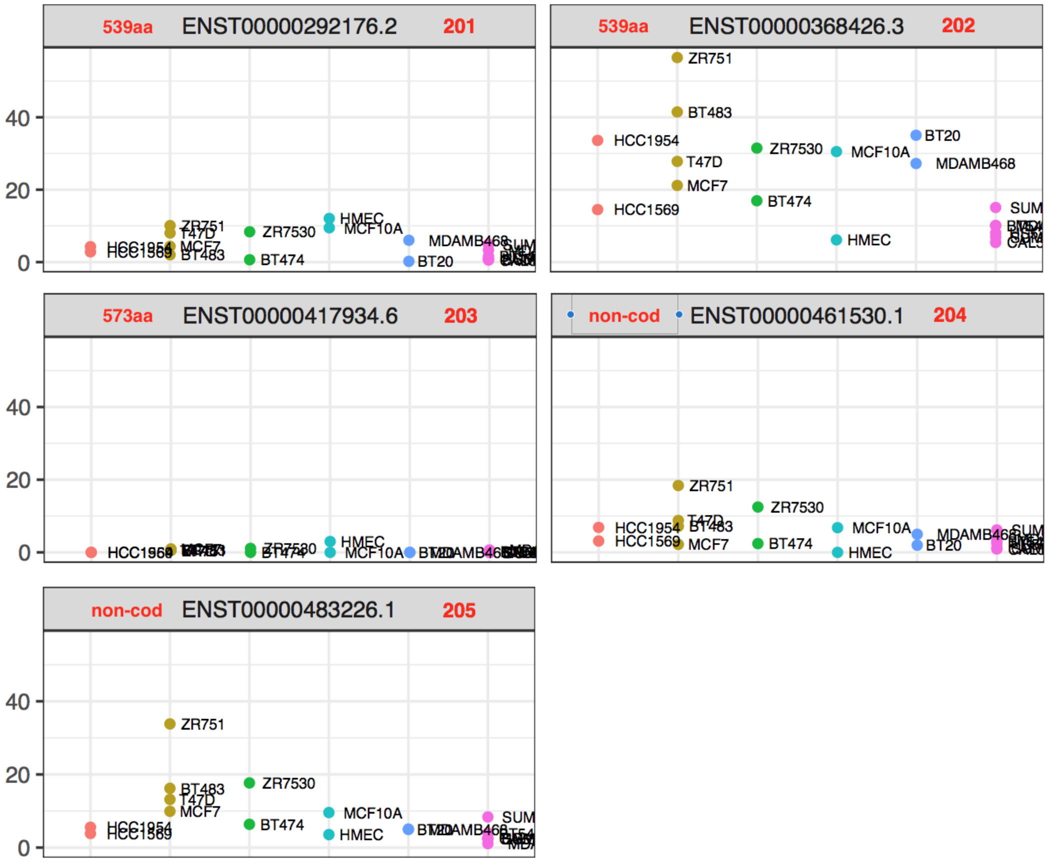

### Supplementary Figure 7

Supplementary Figure 7

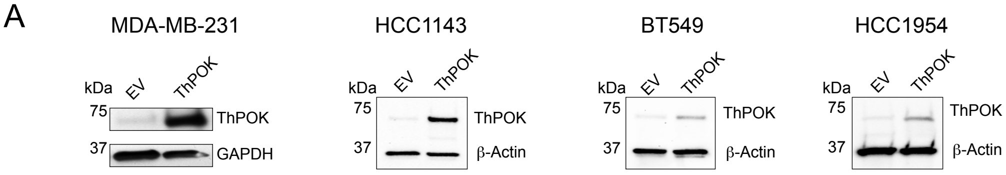


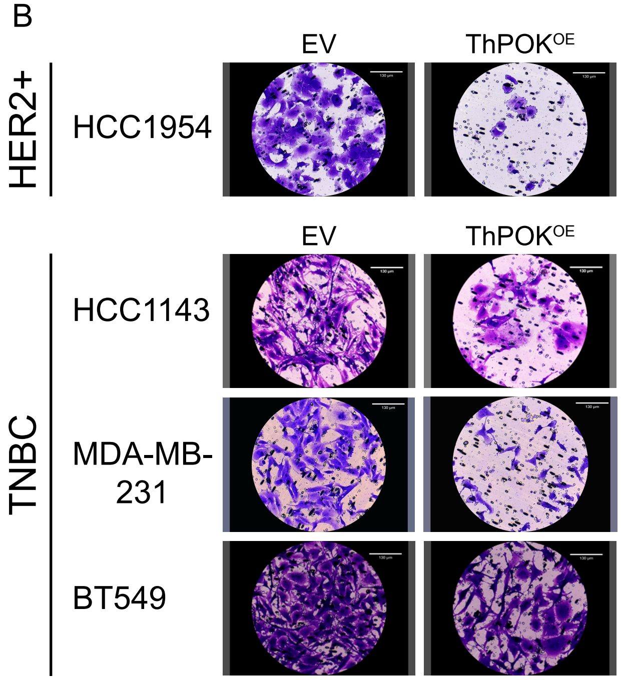

### Supplementary Figure 8

#### Supplementary Figure 8

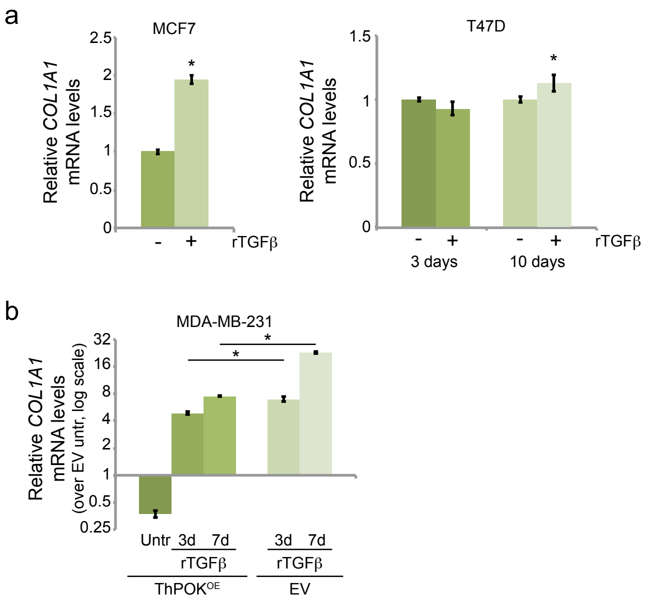
