## Supplementary material for "Super-enhancer profiling reveals ThPOK/ZBTB7B, a CD4^+^ cell lineage commitment factor, as a master regulator that restricts breast cancer cells to a luminal non-migratory phenotype"

### SUPPLEMENTARY METHODS

Unless otherwise indicated, for all data processing we use the statistical language R (1), under the RStudio interphase.

#### Methods for Figure 1 and Supplementary Figure 1

**ChIP-seq data processing and SE analysis**

After read alignment and peak calling, all ChIP-seq samples were subjected to a quality control step. Fingerprint curves of all samples were generated for each dataset using the *plotFingerprint* tool from the *deeptools* suite (2), randomly sampling 100000 500bp-long regions across the genome, with all other parameters as default. Cell lines that showed low quality fingerprints were discarded and, in cases where multiple datasets were available for the same cell line, we used this analysis in order to keep the best one. The final set of ChIP-Seq samples used is indicated in **Supplementary Table 1**.

The reads (*bam*) and peaks (*bed*) files corresponding to the selected datasets were used as inputs to call super-enhancers (SE) for each individual cell line using the ROSE2 algorithm (3, 4) with default options (as described in https://bitbucket.org/young_computation/rose). To compare SE presence between cell lines or subtypes, all overlapping and/or immediately adjacent regions from different cell lines were merged into a single SE set using *bedtools merge*.

H3K27ac signal files (in *bigwig* format) were obtained for each cell line using the *deeptools* (2) *BamCompare* tool. Since the data was single-end, theoretical fragments were generated by extending the reads from both IP and Input files according to the average fragment size on each ChIP sample, obtained from MACS. The signal was calculated using the library-size normalized number of fragments from the IP sample that fall on 25bp-bins covering the whole genome, subtracting the corresponding normalized fragment counts from the Input sample. All negative values were converted to zeroes, to diminish the effect of dissimilar Input qualities between the different datasets.

The total H3K27ac signal on each region of the unique SE set was quantified from the *bigwig* files using *deeptools* (2) *multiBigwigSummary* BED-file tool. This matrix was $\log_{10}$-transformed, scaled and centered by cell line. Next, a principal component analysis (PCA) was performed (**Fig. A**) and pairwise similarity between cell lines was assessed using the Spearman correlation coefficient (**Fig. B**). This exploratory analysis did not identify any cell line as outlier.

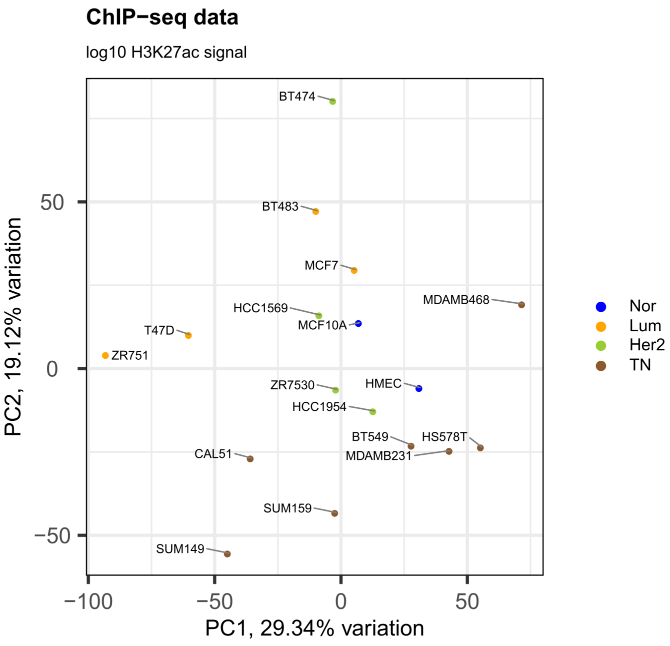


**Figure A**. Principal component analysis of the selected ChIP-seq signal files based on the SE H3K27ac signal, centered and scaled. Samples are colored by subtype.

The heatmaps on **Supplementary Figure 1** were generated with a subset of SE using the scaled and centered H3K27ac signal. First, a filter on the scaled median signal of at least -1 was applied and then the 200 most variable regions were selected. Plots were generated using the *heatmap.2* function from the *gplots* R (5) package, with and without hierarchical clustering of the cell lines.

The SE circo plot was generated using the *circlize* R package.

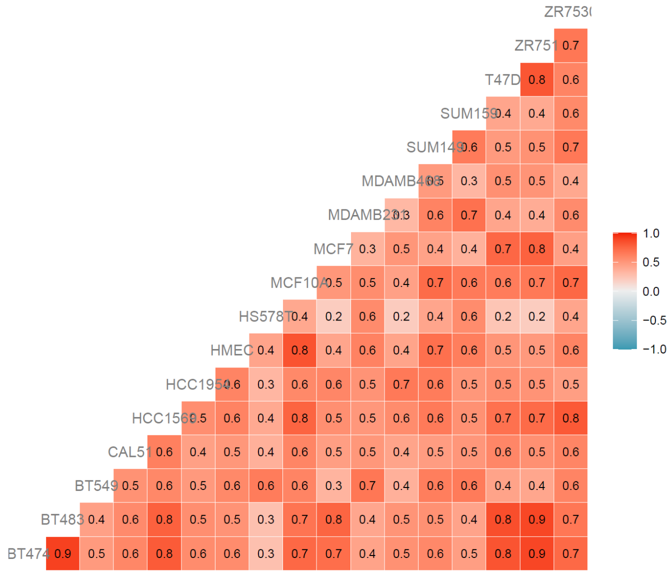


**Figure B.** Spearman correlation analysis of the selected ChIP-seq signal files based on the SE H3K27ac total signal.

**Functional enrichment analysis of SE-associated genes**

Genomic Regions Enrichment of Annotations Tool (GREAT) (7) through the *rGREAT* R (8) package was used to assess the enrichment of SE-associated genes into Gene Ontology (GO) - Biological Processes terms. The gene regulatory domain was defined using the "basal plus extension" rule, setting a distance of 10Kbp upstream and downstream of the TSS to define the basal domain plus a 1Mbp extension. The set of interest and the background set used in each case is described in the main text.

Only those terms with a Benjamini-Hochberg adjusted *p*-value lower than 0.05 and a fold-enrichment ≥ 2 were kept. For visualization purposes, the resulting terms were reduced by similarity of their associated genes using the *rrvgo* R (9) package. The "Rel" similarity measure was selected, with a 0.9 similarity threshold for the analysis on all SE and a 0.7 similarity threshold for the analysis of specific SE sets. The treemap plots were generated using the treemapPlot function from the package on the resulting table of reduced terms.

Differential activity analysis of master regulators

The Virtual Inference of Protein activity by Enriched Regulon analysis (VIPER) method (10) was used to assess differential activity of key transcriptional regulators in specific breast cancer subtypes. VIPER infers the activity of key nodes in a gene regulatory network (termed "master regulators") from the expression levels of their targets. It requires expression data and a regulatory network, including the mode of regulation of their components, as inputs. Expression levels for all genes were taken from RNA-seq data for the different breast cancer cell lines (published data from (11), GEO: GSE73526). A previously built breast cancer regulatory network ("regulonbrca") was used, containing master regulators and their targets (regulons), available through the *aracne.network* R (12) package. RNA-seq read counts were normalized to *tpm* and a graphical exploration of the data was made to ensure that the distribution of gene expression signals was similar between cell lines (**Fig. C**). Also, gene IDs were converted from Ensembl to Entrez format to match those of the “regulonbrca” network. In this process, genes with missing HGNC or Entrez aliases were lost.

A multiple-sample VIPER (msVIPER) analysis was performed for the comparison of each BRCA subtype against all the other samples. Briefly, for each contrast and each regulon, the program calculates and ranks a Gene Expression Signature (GES) to obtain a normalized enrichment score (NES) based on the position shift on the mode of regulation rank. The GES was obtained from the RNA-seq data, after doing a two-group Student's *t*-test and normalizing the *t*-statistics to Z-scores. Enrichment significance was assessed by comparison with a null model, consisting in 1000 uniform and random permutations of the sample labels in the original data. A positive/negative NES is indicative of an over/underactivation of a master regulator on its subtype. Because there is a correlation between the significance of the master regulator differential activity (p-value) and the absolute NES values (**Fig. D**), a threshold of 2 on the absolute NES value was used to select those master regulators with a meaningful change in their activities.

The final set of differentially active master regulators shown in **Figure 1** resulted from keeping those master regulators overlapping any SE from the common SE set.

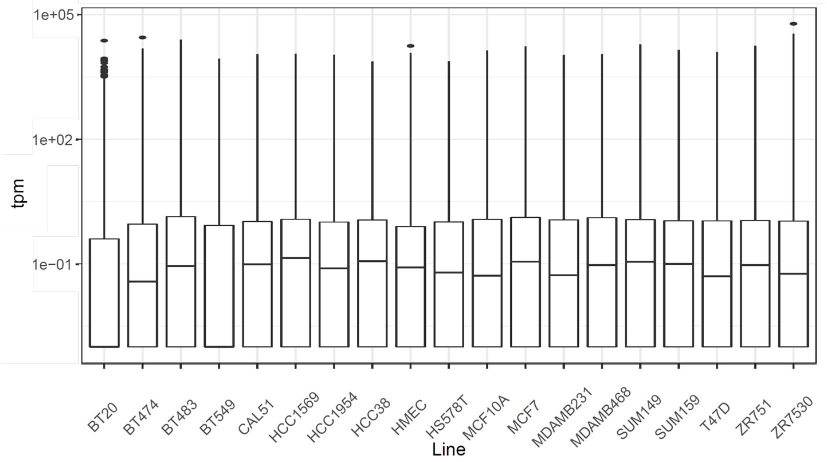


**Figure C.** Gene expression signal (tpm) distribution of all lines analyzed.

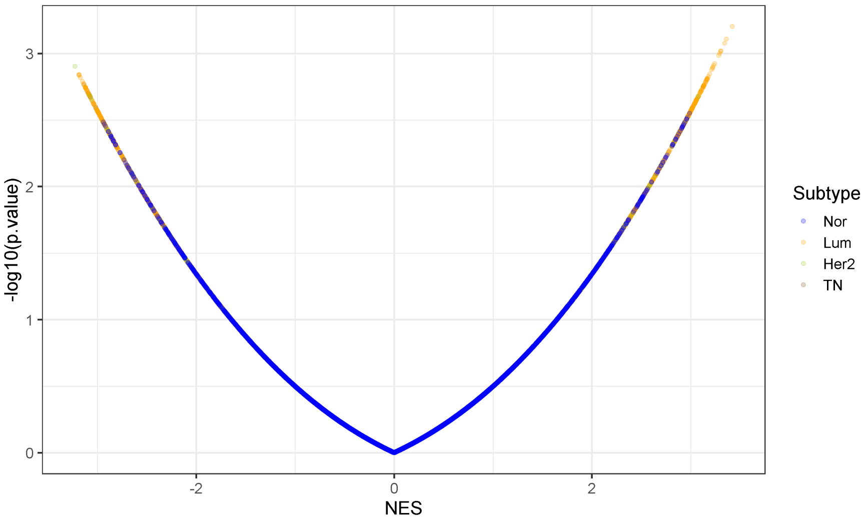


**Figure D.** Volcano plot of the NES values obtained by the VIPER analysis, colored by subtype.

### Methods for Supplementary figure 3

#### TCGA Gene Expression and DNA Methylation Data

TCGA normalized RNA-sequencing FPKM-UQ expression values and normalized DNA methylation beta values, as well as corresponding sample metadata were downloaded using TCGAbiolinks package in R (13–15) . Additional clinical data for ER, PR and HER2 status by immunohistochemistry (IHC) annotation was downloaded using the cgdsr package in R (16) using the Breast Invasive Carcinoma (TCGA, Firehose Legacy) annotation file. IHC receptor status groups were defined as either HR+/- and HER2+/- based on ER, PR and HER2 status by IHC. HR+ includes ER+ PR+, ER+ PR-, ER- PR+, ER+, PR+; HER2+ and HER2- excludes Her2equivocal. PAM50 intrinsic molecular subtypes are annotated as PAM50 Normal, LumA, LumB, HER2 or Basal. Analysis was restricted to samples from women with either PAM50 intrinsic molecular subtype or receptor status by IHC annotation.

Distribution of ThPOK log2(FPKM-UQ+1) expression values in breast cancer tissue were assessed across matched normal (Normal n=112) and PAM50 subtypes (PAM50 Normal n=40, PAM50 LumA n=566, LumB n=207, HER2 n=82, Basal n=194); or IHC receptor status (HR+ HER2+ n=123, HR+ HER2- n=444, HR- HER2+ n=37, HR- HER2- n=115). Non-parametric Kruskal-Wallis test was performed to compare ThPOK expression across groups. Wilcoxon test was then performed for post-hoc pairwise comparisons between groups; *p*-values were adjusted for multiple-testing across all pairwise comparisons using Benjamini-Hochberg (BH) method.

Distribution of ThPOK methylation beta values in breast cancer tissue were assessed across matched normal (Normal n=97) and PAM50 subtypes (PAM50 Normal n=34, PAM50 LumA n=422, LumB n=142, HER2 n=46, Basal n=137). Non-parametric Kruskal-Wallis test was performed to compare ThPOK methylation across groups; p-values were adjusted (BH) for multiple-testing across CpG probes. Wilcoxon test was then performed for post-hoc pairwise comparisons between groups; p-values were adjusted (BH) for multiple-testing across all pairwise comparison and across CpG probes.

### SUPPLEMENTARY FIGURE LEGENDS

**Supplementary Fig. 1**: **Characterization of the consolidated breast cancer cell lines SE set. a)** Heatmaps of the scaled H3K27ac signal (see Supplementary Methods) corresponding to the 200 most variable regions from the set of 5917 SEs across the 17 cell lines. The cell lines were subjected to hierarchical clustering, and the reported subtype is shown in the upper color bar. **b)** Idem to **a**, but the lines are grouped according to its reported subtype. Note that the unsupervised clustering of the cell lines based on this 200 most variable SEs resembles the subtype classification. **c)** Significant GO Biological Process terms found enriched for genes associated to all SEs when performing GREAT analysis. Similar terms (those sharing similar gene sets) are grouped in squares of the same color, and a representative term is shown superposed in white. The full list of terms is shown in **Suppl. Table 3d)**. Individual SE calls from ROSE2 for each of the 17 cell lines around the *ZBTB7B locus*, grouped and colored by subtype. The box highlights the SE in the region overlapping the *ZBTB7B* gene. **e)** Idem to **main Figure 1**, but only the SE region overlapping *ZBTB7B* gene is shown.

**Supplementary Fig. 2: Super-enhancers disruption by BRD4 inhibition leads to decreased *ThPOK* expression. a)** *ThPOK* (left panel) and *ACTB* (right panel) mRNA levels measured by qRT-PCR in luminal (T47D, MCF7) and HER2+ (BT474) cells after 6h treatment with BRD4 inhibitors iBET and JQ1 with the indicated concentrations. **b)** *ThPOK* (left panel) and *ACTB* (right panel) mRNA levels measured by qRT-PCR in TNBC (MDA-MB-231, HCC1143) ThPOK overexpressing cells after 6h treatment with BRD4 inhibitors iBET and JQ1 with the indicated concentrations. For **a-b**, all expression values are depicted relative to the mean of the corresponding controls. Error bars represent the SD of ≥3 biological replicates. NS: non-significant; *: $p < 0.05$; **: $p < 0.005$; ***: $p < 0.0005$; 2-samples *t*-test.

**Supplementary Fig. 3: *ThPOK* gene expression and promoter methylation by breast cancer subtype. a-b)** *ThPOK* expression in TCGA breast cancer subtypes. Boxplots showing distribution of RNA-seq log2 FPKM across **a)** matched normal and PAM50 intrinsic molecular subtypes (Kruskal-Wallis p-value < $3.5\text{x}10^{-34}$); and **b)** matched normal and receptor status by immunohistochemistry (Kruskal-Wallis p-value < $1.32\text{x}10^{-25}$). Post-hoc Wilcoxon p-value significance levels adjusted for multiple testing across all pairwise comparisons are annotated (see below). **c)** *ThPOK* genomic coordinates in UCSC Genome Browser with location of CpG islands and Infinium 450K probes annotated. **d-e)** DNA methylation levels at CpG sites mapping to *ThPOK* across matched normal and PAM subtypes. **d)** Median +/- median absolute deviation of methylation beta values at annotated gene regions: TSS1500, TSS200, 5'UTG, gene body and 3'UTR with Kruskal-Wallis *p*-value significance levels adjusted for multiple testing across CpG probes. **e)** Boxplots showing distribution of methylation beta values with post-hoc Wilcoxon *p*-value significance levels adjusted for multiple testing across all pairwise comparisons and across CpG probes. Significance levels are annotated as follows: adjusted p-value < 0.05 (*), < 0.01 (**), < 0.001 (***), < 0.0001 (****).

**Supplementary Fig. 4: Pan-cancer mRNA and protein ThPOK expression levels. a)** ThPOK protein expression depicted as percentage of patients whose tissues were scored positive for ThPOK staining by IHC using two different antibodies. Patient samples were scored positive when moderate to strong nuclear staining was observed. Two cores were sampled from each individual and protein expression is annotated in tumor cells. For more details refer to: https://www.proteinatlas.org/about/assays+annotation#ih. **b)** Box plot showing the distribution of *ThPOK* mRNA levels expressed in FPKM obtained from TCGA RNA-seq data in 17 cancer types. Box plots shown as median and 25$^{th}$ and 75$^{th}$ percentiles.

<u>Supplementary Fig. 5</u>: ***ThPOK* knockdown leads to morphological as well as transcriptional changes associated with an increase in mesenchymal and stemness markers. a)** ThPOK levels measured by western-blot in luminal (T47D, MCF7 and ZR75.1) and HER2+ (BT474) cells expressing 3 different shRNA against *ThPOK* (kd#1-3) or a control shRNA control (Scramble, Scr). β-Actin was used as loading control. **b)** Morphological changes elicited by ThPOK deficiency images taken at 10X magnification. Scale: each side of the square is 400μm. **c)** CD44 expression in luminal (T47D, MCF7 and ZR75.1) and HER2+ (BT474) cells deficient for ThPOK (kd#1-3) measured by flow cytometry and plotted as the mean fluorescence intensity ratio over the scr-expressing cells. **d)** RNA-based stemness score anti-correlates with ThPOK mRNA levels in TCGA breat tumors' genomic data. Welch's t-test p = 4.169x10$^{-8}$. **e)** Increased migration in luminal ThPOK deficient cells (kd#1-3) measured by Boyden chamber assay (bar 190μm).

<u>Supplementary Fig. 6</u>: **Evaluation of *ThPOK* variants expression in breast cancer cell lines. a)** Schematic of primer design to discern between the expression of *ThPOK* variants #3 and #5, and PCR amplification of *ThPOK* from cDNA of various BCCs to determine whether these cells express variant #3 (PCR product should be 450bp) or #5 (PCR product should be 208bp). **b)** Dot plots showing the isoform-level expression per line for the five different *ZBTB7B* mRNAs annotated in Ensembl. Expression values are based in the RNA-seq data and informed in TPM.

<u>Supplementary Fig. 7</u>: **ThPOK overexpression reduces migration of TNBC and HER2+ cells. a)** ThPOK levels measured by western-blot in TNBC (MDA-MB-231, HCC1143, BT549) and HER2+ (HCC1954) overexpressing ThPOK or an empty vector (EV) control. β-Actin and GAPDH were used as loading control. **b)** Decreased migration in TNBC and HER2+ ThPOK overexpressing cells or EV control, measured by Boyden chamber assay (bar 190μm).

<u>Supplementary Fig. 8</u>: **ThPOK and TGFβ have opposing roles in regulating the expression of ECM genes. a)** *FN1* relative mRNA levels measured by qRT-PCR in MCF7 cells after 3 days treatment with recombinant human TGFβ□□□TGFβ), * p=1.33x10$^{-9}$, and T47D cells after 3 and 10 days treatment with rTGFβ□□*p=5.15x10$^{-6}$. **b)** *COL1A1* relative mRNA levels measured by qRT-PCR in MCF7 cells after 3 days treatment with recombinant

human TGFβ□□□TGFβ), * $p=1.39x10^{-7}$, and T47D cells after 3 and 10 days treatment with rTGFβ□□*$p=0.00991$. **c)** *COL1A1* relative mRNA levels measured by qRT-PCR in MDA-MB-231 cells with $ThPOK^{OE}$ or EV, untreated (Untr) or treated with rTGFβ for 3 (*$p=3.51x10^{-5}$) or 7 days (*$p=3.11x10^{-10}$).
